# Altered calcium responses and antioxidant properties in Friedreich’s ataxia-like cerebellar astrocytes

**DOI:** 10.1101/2024.07.19.604129

**Authors:** Chiara Marullo, Laura Croci, Iris Giupponi, Claudia Rivoletti, Sofia Zuffetti, Barbara Bettegazzi, Filippo Casoni, Ottavio Cremona, Gian Giacomo Consalez, Franca Codazzi

**Affiliations:** Division of Neuroscience, IRCCS San Raffaele Scientific Institute, Milan, Italy; Faculty of Medicine and Surgery, Università Vita-Salute San Raffaele, Milan, Italy

## Abstract

Friedreich’s ataxia (FRDA) is a neurodegenerative disorder characterized by severe neurological signs affecting both the peripheral and central nervous system, caused by reduced levels of the frataxin protein (FXN). While several studies highlight cellular dysfunctions in neurons and various other cell types, there is limited information on the effects of FXN depletion in astrocytes and on the potential non-cell autonomous mechanisms affecting neurons in FRDA. In this study, we generated a model of FRDA cerebellar astrocytes to unveil phenotypic alterations that might contribute to cerebellar atrophy and the degeneration of glutamatergic neurons observed in cerebellar dentate nuclei. We treated primary cerebellar astrocytes with an RNA interference-based approach, to achieve a reduction of FXN comparable to that observed in patients. These FRDA-like astrocytes display some typical features of the disease, such as an increase of oxidative stress, as well as specific functional alterations. Notably, cerebellar astrocytes deplete their reduced glutathione content, becoming more susceptible to oxidative insults. Moreover, FRDA-like astrocytes exhibit alterations of calcium homeostasis, with a reduction in calcium content in the intracellular stores and a corresponding change of calcium responses to purinergic stimuli. Our findings shed light on cellular changes caused by FXN downregulation in cerebellar astrocytes, which can interfere with their physiological and complex interaction with neurons. The potentially impaired ability to provide neuronal cells with glutathione or to release neuromodulators and bioactive molecules in a calcium-dependent manner could impact neuronal function and contribute to neurodegeneration.

## INTRODUCTION

Friedreich’s ataxia (FRDA), a severe progressive neurodegenerative disorder, is the most common inherited autosomal recessive ataxia, affecting approximately 1 in every 50,000 individuals. In most cases, FRDA is caused by biallelic expansion of a naturally occurring GAA repeat in the first intron of the *FXN* gene, resulting in a decrease in its transcription (Campuzano et al., 1996; Keita et al., 2022; Vicente-Acosta et al., 2022). The onset of the disease is observed between the first and the second decade of life and it is inversely correlated with the number of GAA repeats in the *FXN*. Symptoms are progressive and result in severe disability within 10-15 years. Premature death most often occurs because of severe cardiomyopathy (Koeppen, 2011).

FRDA is primarily caused by the deficiency of frataxin (FXN), a ubiquitously expressed mitochondrial protein involved in the synthesis of iron–sulfur clusters, by facilitating their introduction to enzymes containing this prosthetic group (among others, enzymes involved in oxidative phosphorylation and Krebs Cycle). Consequently, FXN deficient cells display a dysregulation of cellular iron metabolism, an increase of reactive oxygen species (ROS) production and, more broadly, of oxidative stress, besides a decreased ATP production and cellular dysfunction (Martelli and Puccio, 2014; Pastore and Puccio, 2013).

While FXN depletion is ubiquitous in FRDA tissues, the cells most clinically affected include large sensory neurons of the dorsal root ganglia (DRG), manifesting degeneration of ascending dorsal columns, dentate nucleus of the cerebellum and other neurons within the retina and the brain. Cardiomyocytes and pancreatic islet cells are also affected. These features contribute to a multisystemic disorder characterized by neurological symptoms such as slowly progressive ataxia of gait and limbs, spasticity, loss of proprioceptive sensation and deep tendon reflexes, dysarthria, visual dysfunction, and hearing loss. Additionally, patients may experience progressive hypertrophic cardiomyopathy, musculoskeletal features, and an elevated risk of diabetes (Harding et al., 2020; Lynch et al., 2021; Pandolfo, 2009).

While the neurodegenerative features of FRDA primarily stem from pathophysiological changes in neurons throughout the peripheral and central nervous systems, FXN depletion may also impact non-neuronal cells (Koeppen, 2011), potentially contributing to the disease’s pathogenesis. Among these cells, astrocytes stand out as the most abundant glial cell type in the nervous system (Sofroniew and Vinters, 2010) and have been implicated in various neurodegenerative disorders, including Huntington’s disease (HD), amyotrophic lateral sclerosis (ALS), multiple sclerosis (MS), Alzheimer’s disease (AD), and Parkinson’s disease (PD) (Sofroniew and Vinters, 2010; Stoklund Dittlau and Freude, 2024; Yuan et al., 2023). Astrocytes are widely recognized for their key contributions to neuronal homeostasis. They provide essential nutrients and metabolic support to neighboring neurons, while also playing a key role in buffering metabolic waste, potassium, and hydrogen ions along their complex processes. Moreover, astrocytes are involved in the handling of neurotransmitters and neuromodulators, as they can uptake and release these signaling molecules, modulating synaptic activity over short and long distances (Navarrete and Araque, 2014). In particular, the ability of astrocytes to remove the excess of glutamate from synaptic clefts plays a crucial role in preventing excitotoxicity, a process implicated in several neurological disorders (Belanger and Magistretti, 2009). Moreover, during inflammatory and oxidative conditions, typical of FRDA, astrocytes undergo an activation process that can have either detrimental or neuroprotective effects. On one hand, activated astrocytes may alter their neuronal support function and release toxic factors, contributing to neuronal death. On the other hand, it is possible that activated astrocytes increase their antioxidant defenses and enhance their ability to handle iron, thus exerting a neuroprotective role (Sofroniew and Vinters, 2010) (Macco et al., 2013; Pelizzoni et al., 2013). Of note, GFAP, a marker of astrocyte activation, has also been detected in the plasma of FRDA patients (Zeitlberger et al., 2018).

Interestingly, selective deletion of *FXN* gene in glial cells of Drosophila, generated FRDA-like symptoms comparable to those of the whole-body knockout flies (Navarro et al., 2010). Furthermore, deletion of Fxn during development in cells with active GFAP promoter, predominantly expressed in astrocytes, induced severe ataxia and premature death, primarily impacting the survival of cerebellar astrocytes (Franco et al., 2017). Recent in vitro studies also showed that FXN knockdown in human astrocytes caused several alterations in these cells (e.g. decreased viability and proliferation, mitochondrial dysfunction) and directed the activation process towards pro-inflammatory and neurotoxic phenotype, contributing to neuron degeneration (Loria and Diaz-Nido, 2015; Vicente-Acosta et al., 2022).

However, in the encephalon the major neuropathological changes linked to FRDA are observed in the cerebellum, a crucial brain region for motor control and cognitive function. FRDA cerebellar signs manifest as progressive atrophy of the dentate nucleus, primarily due to degeneration of large glutamatergic neurons (Koeppen et al., 2011; Selvadurai et al., 2018). Despite this, only limited information is available on cerebellar astrocytes, the consequences of FXN depletion on their phenotype and their possible contribution to cerebellar dysfunction. In the present study we established a model of FRDA cerebellar astrocytes (FRDA-like astrocytes) by *FXN* downregulation, using an RNA interference-based approach in primary cultures of mouse cerebellar astrocytes. Our findings reveal significant functional alterations in FRDA astrocytes, including higher mitochondrial ROS production, increased consumption of the antioxidant molecule glutathione, as well as dysregulation of calcium homeostasis. These results shed light on FRDA pathophysiology in the cerebellum and may contribute to the development of new strategies for disease treatment.

## METHODS

### Primary culture of cerebellar astrocytes

Animal handling and experimental procedures were performed in accordance with the EC guidelines (EC Council Directive 86/6091987) and with the Italian legislation on animal experimentation (Decreto L.vo 116/92) and approved by our Institutional Animal Care and Use Committee.

Mouse cerebella (strain C57BL/6N) were harvested at postnatal day 1 (P1) or 2 (P2) and digested with 2,5 mg/ml Trypsin (Sigma-Aldrich, cat. T1005-1G) and 1,5 mg/ml DNase (EMD Millipore Corp., cat. 260913-10MU). Subsequently, cerebella were mechanically dissociated to obtain a single cell suspension and centrifuged at 100 xg. Pellets were resuspended in astrocyte culture medium (MEM Alpha Medium 1X + GlutaMAX TM (Gibco,cat. 32561-029), 10% Fetal Bovine Serum (EuroClone, cat. ECS5000L), 1% Penicillin-Streptomycin (EuroClone, cat. ECB3001D), 33 mM Glucose (D-(+)-Glucose) (Sigma-Aldrich, cat. G5767-500G)), plated on plastic treated with Poly-L-Lysine (100µg/mL, Sigma, cat. P1274) and cultured in an incubator at 37°, 5% CO_2_. After 2-3 passages, cerebellar astrocytes were plated either on plastic or glass coverslips, both treated with Poly-L-Lysine, depending on experimental requirements.

### Production of lentiviruses expressing *Fxn*-specific and scrambled sh-RNAs

To downregulate the *Fxn* gene in cerebellar astrocytes, vectors encoding for either TRCN0000178380 (5’ GACTTGTCTTCATTGGCCTAT 3’), named sh380, or TRCN0000198535 (5’ GAGTTCTTTGAAGACCTCGCA 3’), named sh535, were generated; plkO.1 SHC002 (5’ ATCTCGCTTGGGCGATAGTGC 3’) named scrambled, was used as control. These vectors were engineered in the laboratory starting from pLKO.1-TRC cloning vectors carrying the shRNAs sequence (Sigma-Aldrich) expressed under the control of the hU6 promoter; the puromycin resistance sequence has been replaced with the sequence encoding for GFP.

Lentiviral particles were prepared as described previously (Amendola et al., 2005). Using the calcium-phosphate precipitation method, HEK293T cells were transiently co-transfected with the transfer vectors, the MDLg/pRRE plasmid, the RSV-Rev plasmid, and the MDLg plasmid. After 72 hours of transfection, cell supernatants containing lentiviral particles were collected, filtered and ultracentrifuged. The pellets were resuspended, divided into aliquots and stored at −80°C. To calculate the transduction efficiency, GFP-positive cells and the total number of cells were counted.

Cerebellar astrocytes were plated on the appropriate support and infected once they reached the desired confluence. The experiments were performed 7 days after the infection.

### Western Blotting

Cerebellar astrocytes were scraped on ice with lysis buffer (0.1M EDTA, 2% NP-40, 0,2% SDS, Clap 1:1000 in PBS). Protein extracts were quantified with Pierce ™ BCA Protein Assay kit (ThermoFisher, 23225). Western blot was performed as described in (Guarino et al., 2022). Briefly, 30μg of protein lysate were suspended in Laemmli sample buffer (final concentration: 50 mM Tris–HCl, pH 6.8, 2.5 mM EDTA/Na, 2% SDS, 5% glycerol, 0.2 M dithiothreitol, 0.01% bromophenol blue), denatured for 5 min at 95°C, loaded onto a 12.5% polyacrylamide gel and then transferred onto nitrocellulose membrane. After 1h of incubation at room temperature (RT) with blocking buffer (TBST - 10 mM Tris/HCl, 150 mM NaCl, 0.1% Tween- 20 - containing 5% skimmed powdered milk), membranes were incubated overnight at 4°C with primary antibodies diluted in blocking buffer and, after extensive washing, with horseradish peroxidase-conjugated anti-rabbit, mouse or chicken secondary antibody (Bio-Rad, Hercules, CA, USA). Proteins were revealed by direct acquisition using the Biorad ChemiDoc™MP Imaging system by Super Signal West Chemiluminescent Substrate (ThermoFisher Scientific). For loading controls, membranes were stripped in a stripping buffer (0.2 M glycine, 0.1% SDS, 1% Tween-20, pH 2.2) and re-probed with the appropriate antibody. Quantification was performed with Image Lab ™Software (Biorad) and protein levels normalized against the loading control (alpha-tubulin).

Primary antibodies: alpha-tubulin (1:6000, Merck, cat. T9026); GFP (1:3000, Abcam, cat. ab13970); Frataxin (1:1000, Merck, cat. AB15080); diluted in 5% Milk in TBST.

### RT-qPCR

RNA was extracted from cells using TRIzol^TM^ Plus RNA Purification kit (Invitrogen, cat. 12183555) and quantified using Nanodrop (ThermoFisher Scientific). 1µg of RNA was retro-transcribed with M-MLV Reverse Transcriptase (Invitrogen, cat. 28025013). RT-qPCR was carried out with LightCycler480 (Roche) using LightCycler480 SYBR Green I Master Mix (Roche).

The primers used for transcript quantifications are:

Frataxin forward primer 5’-TCACCATTAAGCTGGGCG, reverse primer 5’- TTCTTCCCGGTCCAGTCATA; β-actin forward primer 5’-CTGTCGAGTCGCGTCCACC, reverse primer 5’-TCGTCATCCATGGCGAACTG.

The experiment was performed on biological triplicate and RNA levels were normalized to the level of transcript coding for β-actin.

### Fluorescence microscopy Setup

A video imaging setup consisting of an Axioskop 2 microscope (Zeiss, Oberkochen, Germany) and a Polychrome IV light source (Till Photonics, GmbH, Martinsried, Germany) was used for single-cell experiments. Fluorescence images were collected by a cooled CCD videocamera (PCO Computer Optics GmbH, Kelheim, Germany). The ‘‘Vision’’ software (Till Photonics) was used to control the acquisition protocol and to perform data analysis (Rosato et al., 2022; Vangelista et al., 2005). This instrument was used for fura-2-based calcium analyses, mBCl-based GSH measurements and acute iron overload experiments.

The automated ArrayScan XTI platform (Thermo Fisher Scientific) was used for reactive oxygen species (ROS) and TMRM-mitochondrial membrane potential analyses. In the “Results” section this instrument will be referred to as high-throughput microscopy.

### Dye loading and treatments

Dye loading and single-cell experiments were performed in Krebs Ringer Hepes buffer (KRH, containing 5 mM KCl, 125 mM NaCl, 2 mM CaCl2, 1.2 mM MgSO4, 1.2 mM KH2PO4, 6 mM glucose, and 20 mM Hepes, pH 7.4). Experiments were performed at RT.

Fluorescent dyes (from Molecular Probes, Thermo Fisher, when not specified) were loaded as described in the dedicated paragraphs.

After dye loading, cells were washed twice with fresh KRH and analyzed in the same buffer. Acute iron overload protocol was performed incubating cells for 2 min with 20 μM pyrithione (an iron ionophore that allows a kinetically controlled Fe^2+^ entry), before the administration of 100 μM Fe^2+^ (as FAS, ferrous ammonium sulfate) for 3 min, followed by several washes with KRH. To monitor Fe^2+^ entry and iron oxidative status, fluorescence quenching and de-quenching of fura-2 was evaluated at excitation wavelength of 356 nm, the calcium-insensitive wavelength in our optical system.

Chronic iron overload was performed by incubating the cells overnight with 100 μM Fe^3+^ (as ferric ammonium citrate, FAC).

### Analysis of reactive oxygen species (ROS)

Astrocytes were plated on a 96-well plate, transduced with lentiviral particles and analyzed using the Array Scan XTI platform (Thermo Fisher Scientific) one week afterward. Cells were loaded with CellROX Orange Reagent (5 μM, 30 min at 37°C), to analyze ROS accumulation in the mitochondria, and with Hoechst 33342 (5 min at RT, 2 μg/ml final concentration) for nuclear staining.

### Analysis of reduced glutathione (GSH) levels

GSH content was measured at single-cell level by the thiol-reactive fluorescent probe monochlorobimane (mBCl; Sigma-Aldrich); mBCl turns fluorescent after conjugation with GSH. In the mBCl assay, the astrocytes are visualized by means of a nucleic acid stain with SYTO™ (1μM, incubated 10 minutes at 37°C); thereafter 50 μM of mBCl was added to KRH buffer at the beginning of the experiments and the kinetics of fluorescent GSH-monochlorobimane adduct formation was analyzed for 20 minutes, until the plateau phase was reached.

### Mitochondrial membrane potential measurements

Astrocytes were plated on a 96-well plate, transduced with lentiviral particles and analyzed using the Array Scan XTI platform one week afterward.

Cells were loaded with Tetramethylrhodamine methyl ester (TMRM; 15 min at 37°C, 25nM final concentration), in presence of 2μM Cyclosporin H (Cayman Chemical); the solution was maintained in the bath during image acquisition.

Subsequently, 4μM trifluoro carbonyl cyanide phenylhydrazone (FCCP) was added in each well and fluorescence intensity was reacquired on the same cells.

### Calcium Measurement

Cerebellar astrocytes were loaded with fura-2 acetoxymethyl ester (AM, Calbiochem; 40 min at 37◦C, 4 μM final concentration). Images were collected by Axioskope-2-based setup with a rate of 1 ratio image every 2s. The “Vision” software (Till Photonics) was used to control the acquisition protocol and to perform data analysis.

### Immunofluorescence

Astrocytes were plated on a 96-well plate, transduced with lentiviral particles and analyzed using the Array Scan XTI platform after immunofluorescence staining. Cerebellar astrocytes were fixed with 4% paraformaldehyde (PFA4%) in PBS for 15 minutes. The fixed cells were incubated with primary antibodies in 10% Goat Serum (GS), 0.1% Triton in PBS overnight at 4° and, subsequently, with anti-rabbit or anti-mouse secondary antibodies (1:1000, Invitrogen), and counterstained with DAPI (D9542, Sigma).

Images were acquired using Axio Observer (Zeiss Axio Observer.Z1 with Hamamatsu EM9100) and ArrayScan microscope. Quantification of cell types was performed with ImageJ-Fiji or ArrayScan softwares.

Primary antibodies: Glial fibrillary acidic protein (GFAP 1:100, DakoCytomation, cat. Z0334); S100 (β-subunit) (1:500, Sigma-Aldrich, cat. S2532).

### Statistical analysis

Data are expressed as mean +/- standard error of the mean (SEM) of at least 3 independent experiments. GraphPad Prism software was used for the choice of optimal statistical tests based on data distribution. The statistical tests for each experiment are reported in the figure legends. Differences yielding a p value ≥ 0.05 were regarded as non-significant.

## RESULTS

### Downregulation of frataxin expression in cerebellar astrocytes transduced with selective sh-RNAs

To characterize the impact of FXN deficiency on cerebellar astrocytes, we initially established primary astrocytic cultures and defined the conditions for transduction, using lentiviral particles encoding two validated shRNAs, TRCN0000178380 and TRCN0000198535 (hereafter referred to as sh380 and sh535, respectively). The pLKO.1 SHC002 vector, hereon referred to as scrambled, was chosen as the control for the nonspecific effect of lentiviral transduction. To visualize the transduced cells, we replaced one of the two antibiotic resistance sequences, the puromycin cassette, with the GFP coding sequence (refer to **Fig. 1A**). The transduction efficiency, quantified by the ArrayScan instrument, was about 70% for the scrambled construct, 87% for sh535 and 77% for sh380 (evaluated by GFP positivity on >10000, >9000 and >8000 cells, respectively; **Fig. 1B**). We assessed the effect of our knockdown approach on frataxin expression, at both protein and transcript levels. Western blot (WB) analysis, performed 7 days after lentiviral transduction, revealed a reduction of FXN levels to approximately 50% with sh535 and 30% with sh380, compared to untransduced (WT) or control (scrambled-transduced) astrocytes (**Fig. 1C** and **D**). The degree of protein downregulation achieved using sh380 matches FXN levels observed in patient tissues (Gottesfeld et al., 2013). Given the lesser efficiency of sh535, all subsequent experiments were conducted using sh380 only, hereon referred to as sh380-transduced or FRDA-like astrocytes. To compare protein and mRNA downregulation, we performed an RT-qPCR on astrocyte mRNAs, demonstrating a sharp decrease in *Fxn* transcript levels (**Fig. 1E**). The observed differences in protein vs mRNA downregulation are likely explained by the long half life (Li et al., 2008) of the FXN protein, which persists even one-week after lentiviral infection. Again, WT and scrambled-transduced astrocytes displayed comparable *Fxn* transcript levels (**Fig. 1E**). Having established an *in vitro* model of FRDA-like cerebellar astrocytes, we evaluated the impact of FXN downregulation on their morphological features. Indeed, previous studies (Pastore et al., 2003) showed cytoskeletal alterations in FRDA patient fibroblasts, characterized by increased glutathionylation of actin filaments and disorganization of microfilaments. In our study, we performed an immunofluorescence analysis using a GFAP antibody. Our results revealed GFAP positivity of about 43% in control astrocytes, and 48% in FRDA-like astrocytes (**Supplementary figure 1A and B)**. However, the expression of S100β, a calcium binding protein abundantly expressed in astrocytes, was scored in approximately 99% of the cells, both scrambled- and sh380-transduced (**Supplementary figure 1C and D)**. The heterogeneity observed in our primary cultures reflects the highly diverse morphologies of cerebellar astrocytes observed both in culture and *in vivo* models (Buffo and Rossi, 2013; Cerrato, 2020). However, cellular phenotypes were not overtly affected by changes in FXN levels and remained comparable in scrambled and FRDA-like astrocytes (**Supplementary Fig. 1 A and C**).

**Figure 1.**
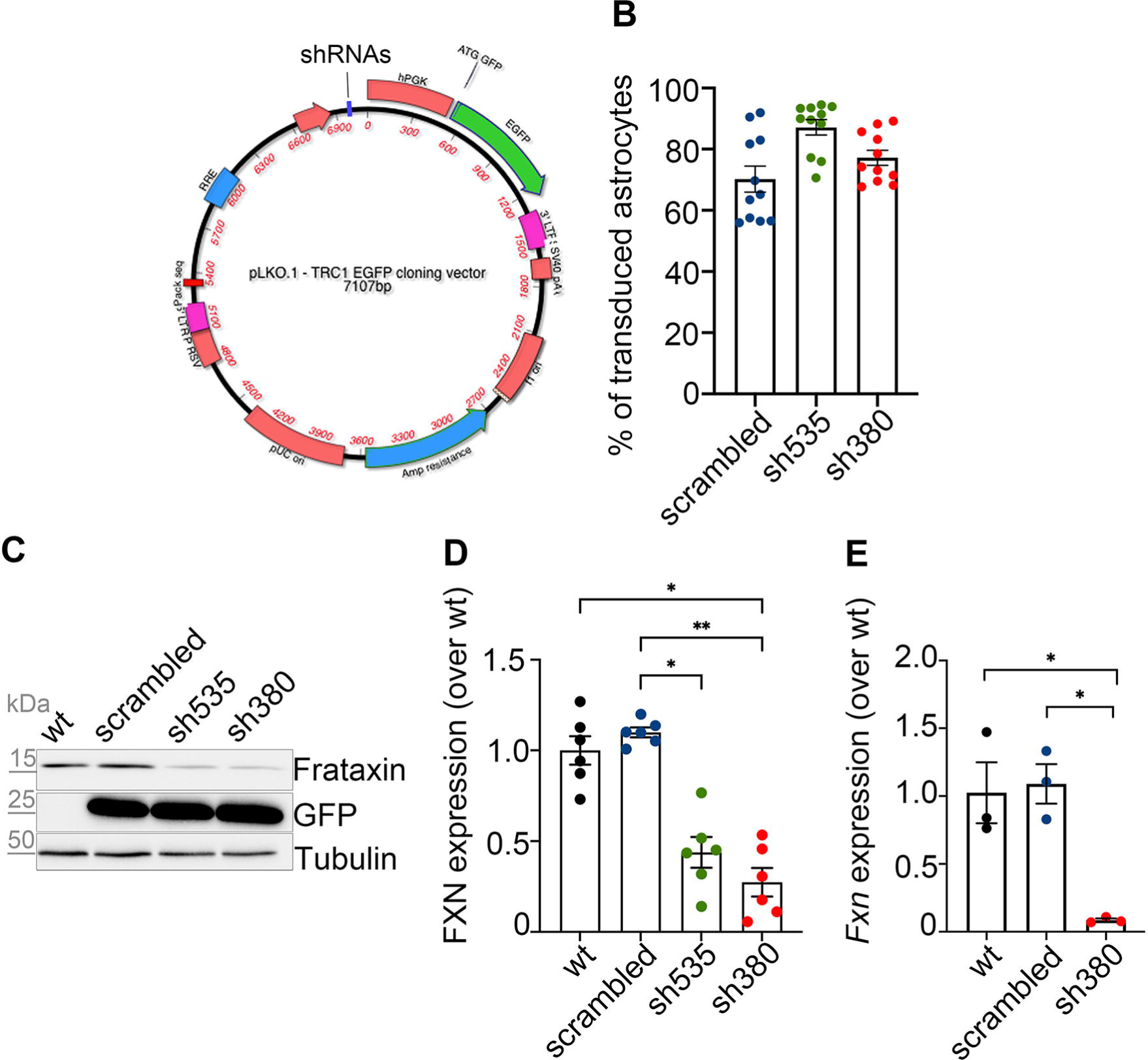
Frataxin knockdown in cerebellar astrocytes. **A**: Representation of the pLKO.1-TRC cloning vector encoding one of three different shRNAs (either scrambled, sh535, or sh380) and the enhanced green fluorescent protein (EGFP). The expression of shRNAs is driven by the hU6 promoter. Ampicillin resistance was used for plasmid selection. **B**: The graph shows the percentage of lentivirally transduced astrocytes assessed by high-throughput microscopy; n=10107 cells for scrambled, 9771 for sh535, and 8263 for sh380, from 5 biological replicates; each dot represents the mean percentage of transduced cells in a single culture well. **C**: Western blot of protein lysates derived from wt, scrambled-, sh535- and sh380-transduced astrocytes, immunostained with antibodies detecting FXN (15kDa). GFP (25kDa) was used to assess the transduction efficiency; ɑ-tubulin (50kDa) was used as a loading control. **D**: FXN protein levels, normalized to ɑ-tubulin levels, in scrambled-, sh535-, and sh380-transduced astrocytes, compared to wt astrocytes. Data are expressed as mean +/- SEM; n=6; Kruskal-Wallis test; *: p<0.05, **: p<0.01. **E**: *Fxn* mRNA levels, normalized to β-actin, in scrambled- and sh380-transduced astrocytes, determined by RT-qPCR compared to wt. Data are expressed as mean +/- SEM; n=3; Brown-Forsythe and Welch ANOVA tests; *p<0.05

### Effects of FXN downregulation on the oxidative status of cerebellar astrocytes

A distinctive hallmark of FRDA is represented by oxidative stress, which has been observed in animal and cellular models, as well as in patient tissues (Lupoli et al., 2018; Pilotto et al., 2024). To assess whether our astrocytic model recapitulates the pathophysiology of FRDA, scrambled- and sh380-transduced astrocytes were loaded with Cell-ROX Orange, a mitochondrial probe whose fluorescence depends on ROS-mediated oxidation. Our results were obtained using high-throughput microscopy to extend the analysis to a large number of cells (Vannocci et al., 2018). Our results showed a significant increase in ROS levels in FRDA-like astrocytes compared to controls (**Fig. 2A**).

**Figure 2.**
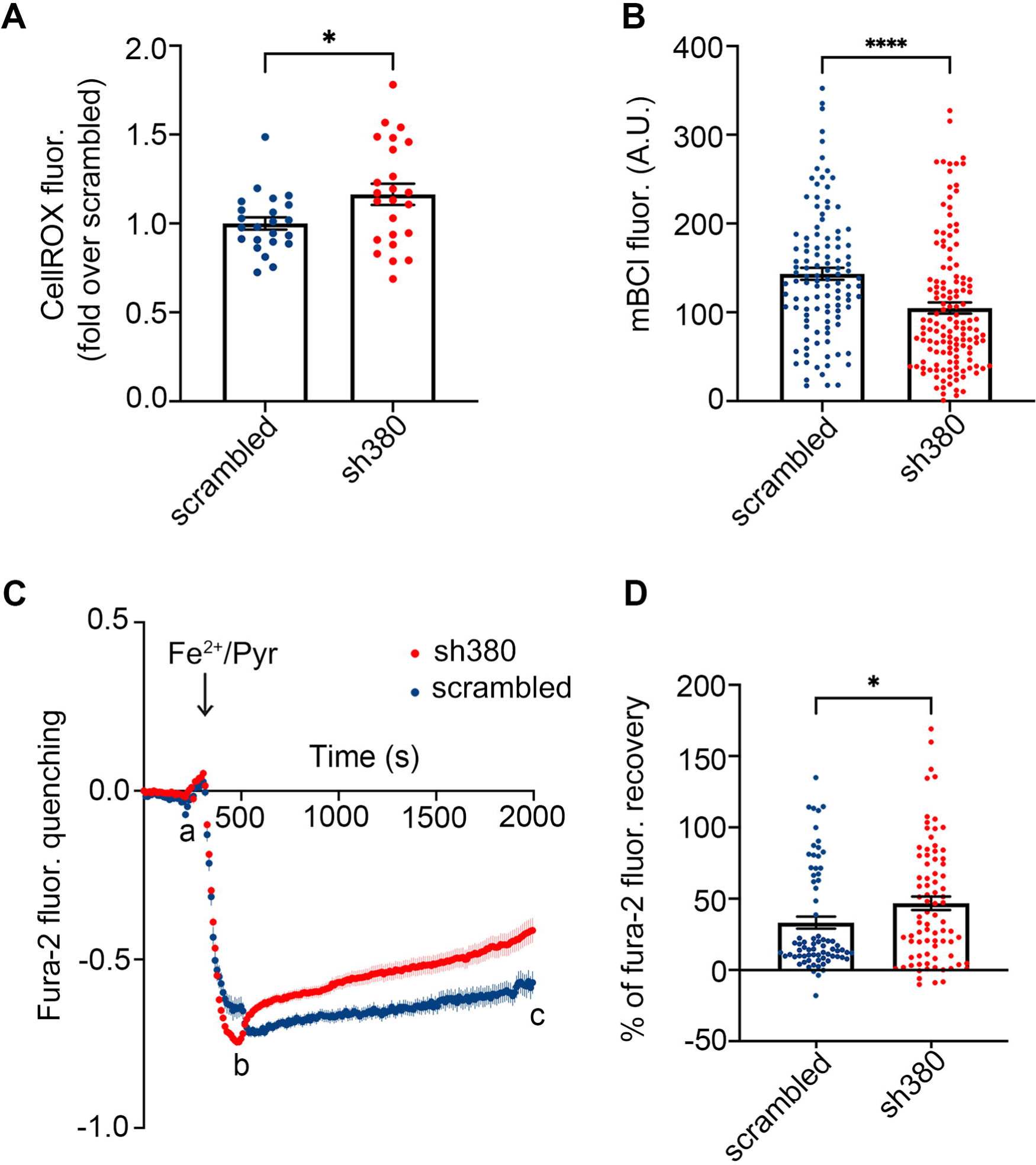
Increased ROS levels and impaired antioxidant defenses in FRDA-like astrocytes. **A**: Quantification of ROS production, analyzed at the single-cell level by high-throughput microscopy, and evaluated by CellROX fluorescence in sh380-compared to scrambled-transduced cells. Each dot represents the average of an individual culture well. Data, from 4 biological replicates, are expressed as mean +/- SEM; n=3822 cells for scrambled, and 2192 for sh380; Mann-Whitney test; *p<0.05 **B**: Measurement of GSH content, estimated in single-cells by mBCl fluorescence, in scrambled and sh380-transduced astrocytes. Each dot represents the fluorescence levels recorded at the plateau phase (20 minutes after mBCl administration) in each individual cell. Data, from 3 biological replicates, are expressed as mean +/- SEM; n=90 cells for scrambled, and 179 for sh380; Mann-Whitney test. ****p<0.0001 **C**: Fura-2 fluorescence kinetics, analyzed at the calcium-insensitive excitation wavelength of 356 nm, upon administration of Fe^2+^ (100μM) in the presence of pyrithione, an iron ionophore; (20 μM), in scrambled- and sh380-transduced astrocytes (blue and red, respectively). Fura-2 basal fluorescence (a) was quenched by Fe^2+^ entry induced by Fe^2+^/Pyr administration (b); the oxidation of Fe^2+^ to Fe^3+^ eventually caused fluorescence recovery (c). Data, from 3 biological replicates, are expressed as mean +/- SEM; n=74 cells for scrambled, 81 for sh380. **D**: Percentage of fura-2 fluorescence recovery in scrambled- and sh380-transduced astrocytes. Data correspond to the ratio of fluorescence recovery to fluorescence quenching [(c-b)/(b-a)]. Each dot represents the fluorescence of a single cell. Data, from 3 biological replicates, are expressed as mean +/- SEM; n=74 cells for scrambled, and 81 for sh380; Mann-Whitney test; ***p<0.001.

As the primary antioxidant defense in glial cells is represented by reduced glutathione (GSH), we assessed whether increased ROS accumulation could interfere with GSH levels. To this end, cerebellar astrocytes were loaded with monochlorobimane (mBCl), a probe that fluoresces upon conjugation with GSH (Bettegazzi et al., 2019). The fluorescence was quantified after 20 minutes, when the reaction reached the plateau phase. In FRDA-like astrocytes the final fluorescence level was significantly lower compared to control astrocytes, indicating GSH consumption, in a likely attempt to counteract the ROS increase caused by FXN deficiency (**Fig. 2B**).

Based on this evidence, we asked whether the decrease in GSH levels scored in FRDA-like astrocytes could impair their antioxidant defenses. To this end, cerebellar astrocytes were loaded with fura-2 and exposed to an acute Fe^2+^ overload (100 μM) in the presence of pyrithione. The unique property of fura-2, whose fluorescence is selectively quenched by Fe^2+^ but not Fe^3+^, makes it possible to measure intracellular iron oxidation and to measure the ability to maintain the intracellular reducing potential (Pelizzoni et al., 2011). In our astrocytes, whereas the acute Fe^2+^ entry caused a similar fluorescence quenching in both controls and FRDA-like astrocytes (**Fig. 2C**), fura-2 fluorescence recovery was significantly higher in FRDA-like astrocytes (**Fig. 2D**), indicating their higher oxidative potential.

### Effects of FXN downregulation on mitochondrial membrane potential

Given the increase in basal ROS levels in sh380-transduced astrocytes, which is often caused by a reduction of the mitochondrial membrane potential (ΔΨ_m_), we investigated this parameter using TMRM, a fluorescent lipophilic dye that accumulates in active mitochondria due to their negative membrane potential. A qualitative analysis of TMRM fluorescence revealed a comparably healthy appearance of mitochondria in FRDA-like astrocytes and in scrambled controls, without signs of mitochondria fragmentation (**Fig. 3A**). Surprisingly, high-throughput microscopy analysis, previously used for ROS measurements, showed that ΔΨ_m_ in FRDA-like astrocytes was slightly higher (although not significantly different) than in control cells (**Fig. 3B**). After the first fluorescence acquisition, the astrocytes were then treated for 10 minutes with FCCP, an uncoupler of the respiratory chain that dissipates the ΔΨ_m_, followed by a second round of acquisition on the same cells. The relative decrease in TMRM fluorescence induced by FCCP was significantly greater in FRDA-like astrocytes, indicating a higher proton gradient than in control astrocytes (**Fig. 3C**). The apparent discrepancy between the untreated and the FCCP-induced ΔΨ_m_ may be attributed to low dynamic range of TMRM fluorescence changes when close to saturation (in untreated cells). This saturation can dampen the differences between control and FRDA-like astrocytes, which are instead visible at fluorescence variations within the linear range.

**Figure 3.**
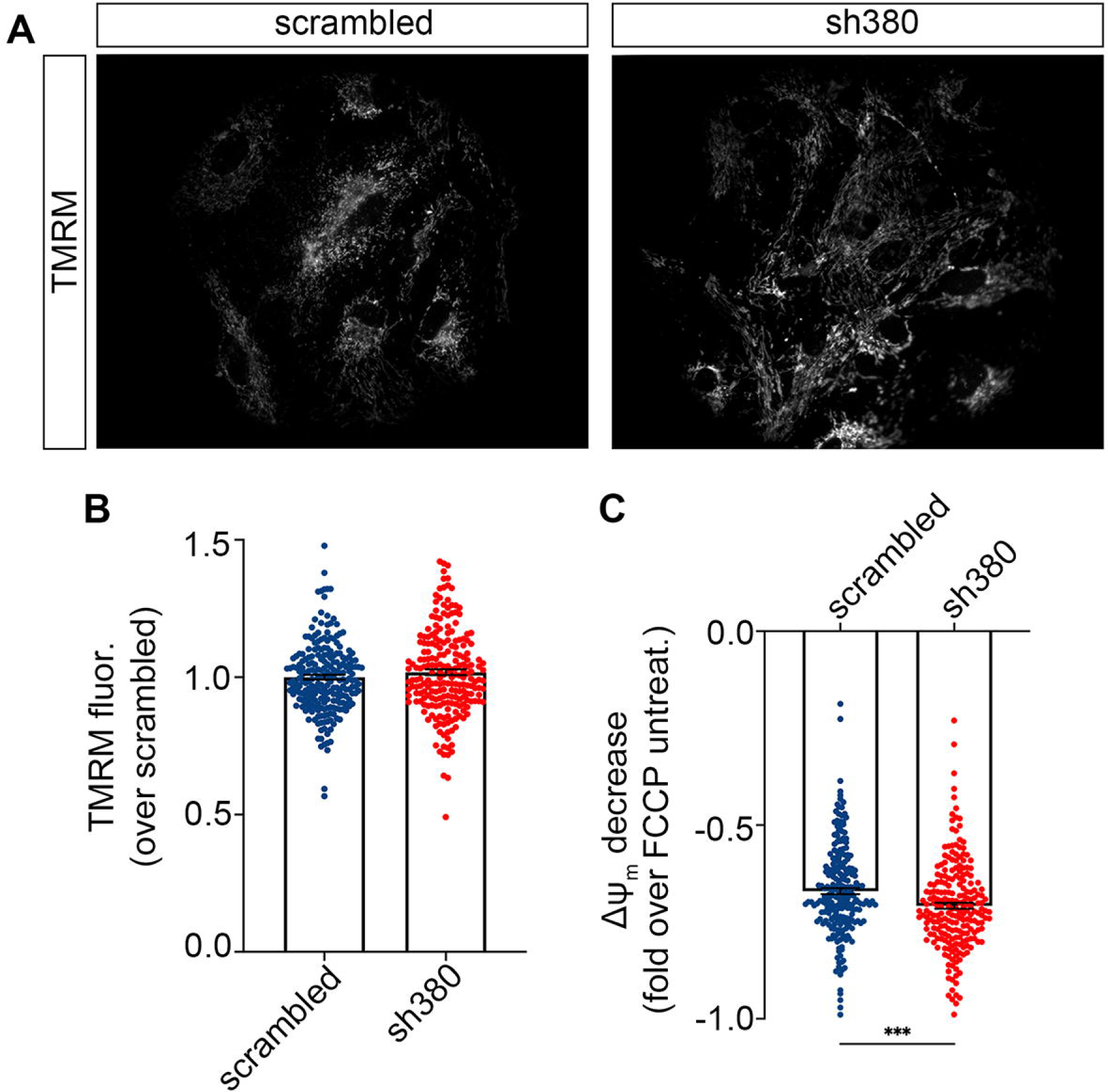
Unaffected mitochondrial membrane potential (ΔΨ_m_) in FRDA-like astrocytes. **A:** Representative images of mitochondria loaded with TMRM, from scrambled- and sh380-transduced astrocytes. **B and C:** Scrambled- and sh380-transduced astrocytes loaded with TMRM were analyzed by high-throughput microscopy. After the first run of acquisition (B), astrocytes were treated with FCCP (4 μM) for 10 minutes and the fluorescence acquisition was repeated on the same cells (C). In C, the fluorescence decrease caused by FCCP treatment was normalized to the untreated condition. Each dot represents the fluorescence of a single cell. Data, from 3 biological replicates, are expressed as mean +/- SEM; n =229 cells for scrambled, and 222 for sh380; Mann-Whitney test ***p<0.001.

### Alterations of calcium homeostasis in FRDA cerebellar astrocytes

Oxidative stress affects several metabolic pathways and cellular functions, with a particular impact on the complex mechanisms underlying Ca^2+^ homeostasis. To analyze calcium metabolism in FRDA-like astrocytes, one week after lentiviral infection, sh380- and scrambled-transduced astrocytes were loaded with fura-2 calcium dye, and analyzed by single-cell calcium imaging. Strikingly, neither astrocyte population responded to glutamate, while all cells showed a robust increase of intracellular Ca^2+^ concentration ([Ca^2+^]_i_) upon acute administration of 100 µM ATP. Surprisingly, the peak of Ca^2+^ responses induced by ATP was significantly reduced in FRDA-like astrocytes compared to scrambled controls (**Fig. 4A**). One of the mechanisms responsible for the amplitude of the Ca^2+^ elevation induced by metabotropic, but also ionotropic, stimuli, is the regulated release of Ca^2+^ from intracellular Ca^2+^ stores. To investigate possible alterations in this pathway, the astrocytes were exposed to thapsigargin, a blocker of SERCA pumps and, consequently, of store refilling. The [Ca^2+^]_i_ increase, resulting from thapsigargin-mediated store depletion, was significantly lower in FRDA-like astrocytes compared to control ones, indicating an impairment of Ca^2+^ storage (**Fig. 4B and C**). Several reports suggest that Ca^2+^ storage and its release from intracellular stores can be affected by the oxidative environment (Ying et al., 2008), which is elevated in FRDA-like astrocytes. To investigate this aspect, we treated the cells for 24 hours with Fe^3+^ (administered as ferric ammonium citrate, FAC, 50 μM), to induce a mild iron overload and increase the oxidative environment. Indeed, under this condition, the elevation of [Ca^2+^]_i_ induced by ATP was even less pronounced in FRDA-like astrocytes than in control cells (**Fig. 4D**).

**Figure 4.**
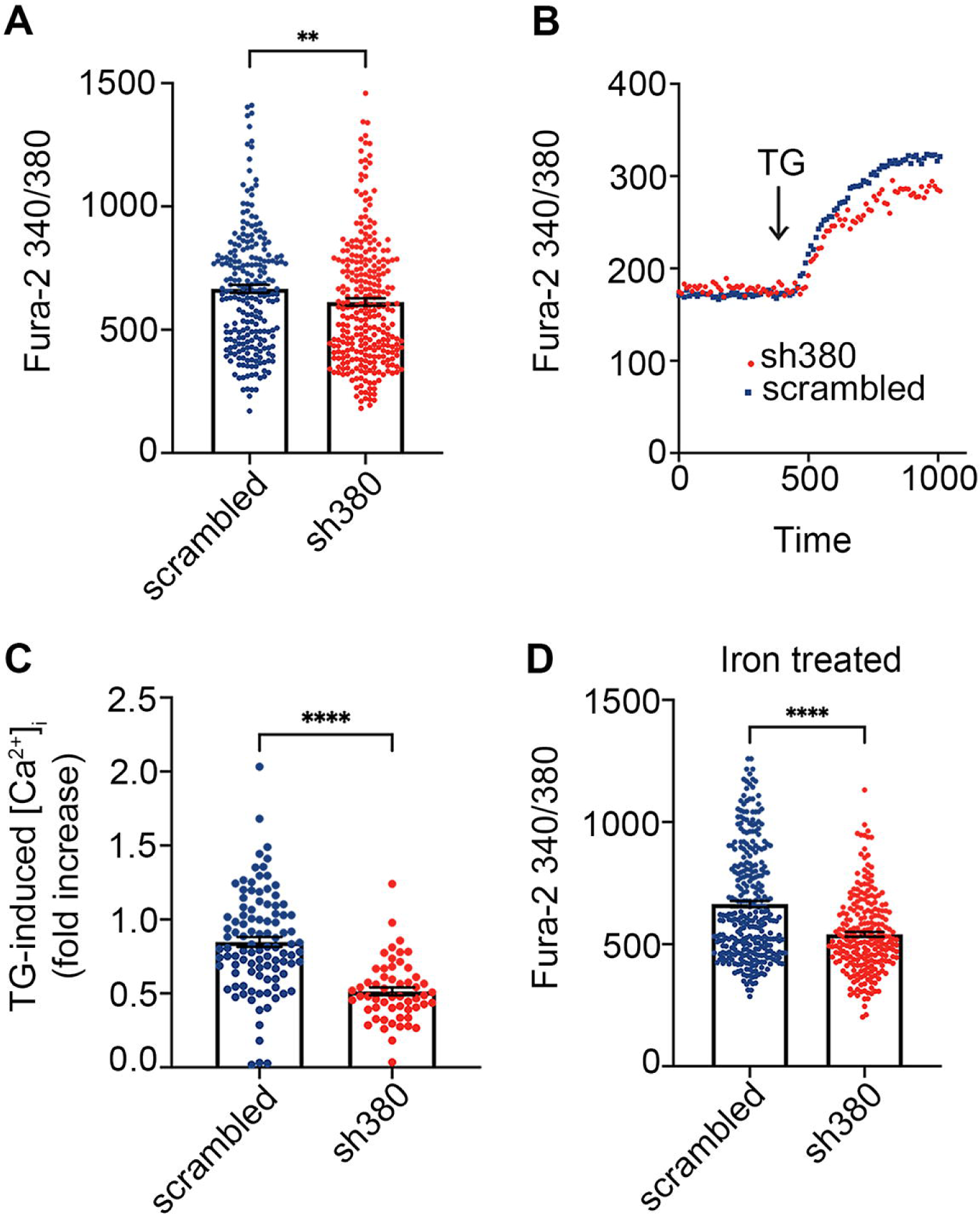
Decreased calcium responses in FRDA-like astrocytes. **A**: Peaks of calcium response following acute ATP administration (100 µM) in scrambled- and sh380-transduced astrocytes. Each dot represents the maximum peak scored in each single cell. Data, from 5 biological replicates, are expressed as mean +/- SEM; n=237 cells for scrambled, and 292 for sh380; Mann Whitney test; **p<0.01 **B and C**: [Ca^2+^]_i_ increase caused by calcium store depletion following thapsigargin (TG) treatment. In B representative kinetics of [Ca^2+^]_i_ elevation upon TG administration (1 µM). In **C**, TG-mediated depletion of calcium stores. Each dot in C represents a single cell. Data, from 3 biological replicates, are expressed as mean +/- SEM; n=101 cells for scrambled and 87 for sh380; Unpaired t test; ****p<0.0001 **D**: Peaks of calcium response following ATP administration (100 µM) in cerebellar astrocytes treated for 24 hours with Fe^3+^ (administered as ferric ammonium citrate, FAC, 50 μM). Each dot represents the maximum fluorescence peak scored in each individual cell. Data, from 5 biological replicates, are expressed as mean +/- SEM; n=313 cells for scrambled, 237 for sh380; Mann Whitney test; ****p<0.0001

## DISCUSSION

For many decades, research on neurodegenerative disorders has focused primarily on the intrinsic mechanisms of neuronal toxicity. However, mounting evidence demonstrates that glial cells actively contribute to pathological processes and disease progression. Astrocytes, whose physiological role in neuronal homeostasis and function is very well established, can lose their supportive properties during a neurotoxic process, and actively contribute to neurotoxicity (Gleichman and Carmichael, 2020; Stoklund Dittlau and Freude, 2024). In FRDA, the potential impact of these non-cell autonomous mechanisms on the disease has only been partially investigated.

In this study, we generated and characterized an in vitro model of FRDA cerebellar astrocytes, to evaluate their vulnerability to FXN knockdown and their potential contribution on FRDA pathogenesis. Indeed, selective ablation of mouse *Fxn* in GFAP-expressing precursor cells causes alterations selectively in cerebellar astrocytes, but not in forebrain astrocytes, leading to severe ataxia and early death (Franco et al., 2017). This finding suggested that cerebellar astrocytes are more vulnerable to frataxin depletion than astrocytes from other brain regions. This specific cerebellar vulnerability could be partially attributed to the unique proteomic profile of astrocytes from the cerebellum, which differ from that of astrocytes in all other brain regions analyzed (Prabhakar et al., 2023).

The cerebellum is characterized by astrocyte heterogeneity, resulting from a complex and still partially unclear process of glial cell proliferation and differentiation. Astroglia originate from radial glia of the ventricular neuroepithelium through direct transformation into Bergmann glia at embryonic stages and by amplification of intermediate progenitors within the cerebellar parenchyma after birth (Cerrato et al., 2018). Based on their morphological features and functional connections, cerebellar astrocytes can be classified in three main groups: the Bergmann glia, the star-shaped astrocytes (also defined velate protoplasmic astrocyte) of the granular layer in the cerebellar cortex, and fibrous astrocytes in the white matter (Buffo and Rossi, 2013; Cerrato, 2020). Our cultures, obtained from cerebella harvested at neonatal stages, likely reflect this heterogeneity, with cells characterized by varying morphologies (stellate, fibroblast-like, elongated) and different expression of the typical astrocytic markers (GFAP and S100β). The low percentage of GFAP-positive cells found in our cultures mirrors similar behavior observed in rat hippocampus (Walz and Lang, 1998) and cerebral cortex (Zhang et al., 2019). Within this framework, no morphological differences seem to be induced by FXN knockdown, likely because the in vitro differentiation program, once initiated, is no longer affected by FXN silencing. GFAP, a typical marker that significantly increases in reactive astrocytes and during astrogliosis, was also found to be elevated in the cerebellar astrocytes where FXN was downregulated at early stages (Franco et al., 2017). However, in our cultures, under pro-inflammatory conditions, GFAP levels sharply decreased compared to resting conditions (unpublished data). Again, this behavior does not depend on FXN depletion, but it should deserve attention when evaluating the effects of inflammatory conditions occurring during FRDA progression (Hayashi et al., 2014; Koeppen et al., 2016).

Having obtained an FRDA-like cerebellar astrocytic model, showing a decrease in FXN levels comparable to that observed in patients, we investigated the functional alterations caused by FXN downregulation. Impaired mitochondrial metabolism and a defective respiratory chain, associated to ROS production, are considered one of the main mechanisms of FRDA pathogenesis (Calabrese et al., 2005; Lupoli et al., 2018). Indeed, a typical hallmark of FRDA is a reduction in ΔΨ_m_, observed in affected neurons as well as in various cell types. However, our FRDA-like astrocytes do not display noticeable morphological changes in mitochondria, nor a significant alteration in ΔΨ_m_. Interestingly, this parameter appears slightly higher, although not significantly different, in sh380-transduced astrocytes compared to scrambled-transduced ones. Accordingly, the effect of FCCP, an uncoupler of the respiratory chain that disrupts the proton gradient, was more pronounced in FRDA-like astrocytes than in the control counterpart. Our results are consistent with data obtained from cerebellar cultures, where the ΔΨ_m_ was significantly decreased in granule cells of FRDA YG8R mice, but not in glial cells, where this parameter was comparable to cultures from control mice (Abeti et al., 2016). This finding suggests a compensatory mechanism in glial cells that maintains normal ΔΨ_m_ under FXN depletion.

With respect to the observed increase in oxidative stress, the higher production of free radicals, typically observed in FXN-deficient cells and associated to oxidative damage in FRDA (Codazzi et al., 2016), depends on the reduction of ΔΨ_m_. Nonetheless, our FRDA-like astrocytes showed higher levels of basal ROS production, despite the unchanged ΔΨ_m_. However, it has been shown that FXN intrinsically increases cellular antioxidant defenses by activating glutathione peroxidase and elevating reduced thiols (Shoichet et al., 2002). Therefore, the downregulation of FXN accounts for higher oxidative stress in FRDA-like astrocytes. It is important to note that ROS generation in FRDA-like astrocytes is not excessively elevated, likely due to their high ability to buffer perturbation of various intra- and extracellular parameters. Indeed, our cerebellar FRDA-like astrocytes consume their GSH content to counteract the basal ROS elevation. However, GSH depletion impairs their ability to maintain the redox cellular balance, as observed under a condition of acute iron overload (Pelizzoni et al., 2011), making the astrocytes more vulnerable to oxidative insults. This condition also impairs the protective role of astrocytes towards neuronal cells, which depend entirely on astrocytes for their GSH content (Aschner, 2000).

Cerebellar FRDA-like astrocytes also display alterations in Ca^2+^ homeostasis that might affect the physiological functions of astrocytes. Astrocytes generate complex Ca^2+^ signals, such as Ca^2+^ oscillations and waves (Codazzi et al., 2001), in response to neurotransmitter spillover from the synaptic cleft and to other extracellular signals. In turn, the spatiotemporal aspects of Ca^2+^ dynamics permit a controlled release of active molecules and neuromodulators, both locally and within long-range neuro-astrocytic networks. Spontaneous and repetitive Ca^2+^ waves have been observed and characterized in Bergmann glia in vivo (e.g. Asemi-Rad et al., 2023; Hoogland and Kuhn, 2010). These calcium signals seem only partially mediated by glutamatergic AMPA receptors, whose Ca^2+^ permeability depends on the expression of the GluR2 subunit. Likewise, the stimulation of purinergic receptors, particularly the metabotropic P2Ys, promotes complex Ca^2+^ responses (Hoogland and Kuhn, 2010). Our data confirm a lack of calcium signals induced by glutamate stimulation (not shown) and consistent responses to ATP. Moreover, we demonstrated that ATP-induced Ca^2+^ responses in FRDA-like astrocytes are lower in comparison to controls and that a lower Ca^2+^ loading of intracellular stores (revealed by thapsigargin-induced store depletion) accounts for Ca^2+^ mishandling in FRDA-like astrocytes. A similar impairment of the ability to load Ca^2+^ in intracellular stores has been observed in cerebellar granule neurons derived from the YG8R FRDA mouse model (Abeti et al., 2018). This altered Ca^2+^ homeostasis can be caused by the higher level of oxidative stress displayed by FRDA-like astrocytes, due to the presence of cysteines in SERCAs that are sensitive to oxidative environments (Ying et al., 2008). Moreover, while it is more common to observe elevated calcium responses under toxic cellular conditions, a similar decrease of Ca^2+^ signals have been observed in other neurodegenerative models. For instance, in cellular models of TDP-43 proteinopathy, a feature of amyotrophic lateral sclerosis (ALS), glutamate-elicited Ca^2+^ responses were significantly lower than in control cells (Pisciottani et al., 2023). Likewise, Müller cells in the retina, maintained in high glucose to mimic diabetic retinopathy, also showed Ca^2+^ responses to ATP stimulation that was reduced compared to cells maintained in normal-glucose medium (Rosato et al., 2022). Similarly, in astrocytes, the activation of P2Y receptors enhances resistance to oxidative stress by a Ca^2+^-dependent increase of mitochondrial metabolism (Wu et al., 2007). Therefore, lower Ca^2+^ responses can impair this protective pathway. Additionally, the release of several active molecules and factors by astrocytes depends on Ca^2+^ elevation, so impaired responses to ATP could affect and alter their physiological interplay with neuronal cells.

## CONCLUSIONS

In this study, we characterized for the first time a set of functional alterations in FRDA-like cerebellar astrocytes. Although FXN downregulation does not seem to cause macroscopic morphological changes, in-depth analyses revealed modification of some cellular parameters that might significantly affect long-term neuronal function and synaptic activity. The reduction of astrocytic GSH might indeed compromise the already low antioxidant capacities of neurons. This might accelerate disease progression, increasing the neurotoxic impact of iron accumulation and oxidative stress. Moreover, the reduced calcium responses to purinergic stimuli might alter the ability of astrocytes to release molecules that actively modulate and regulate synaptic function over long distances, further contributing to neuronal dysfunction. The most overtly affected neurons in the brain are the large glutamatergic neurons of the dentate nuclei, which are not easily co-cultured with FRDA-like astrocytes. Different in vitro models (e.g. organotypic cerebellar slices maintained with medium conditioned by astrocytes) will make it possible to assess how FXN depletion in cerebellar astrocytes might alter their complex interplay with glutamatergic neurons and contribute to neuronal dysfunction. Likewise, selective stereotaxic injection of adeno-associated viral particles into the mouse lateral nucleus (e.g. Asemi-Rad et al., 2023) to achieve cerebellar astrocyte-specific FXN knockdown would provide a useful in vivo model of locally altered astrocyte/neuron interactions and of their impact on the ataxic phenotype.

### Data availability statement

The raw data supporting the conclusions of this article will be made available by the authors, without undue reservation.

## Supporting information

Supplemental Figures Marullo 2024

## Author contributions

CM: investigation, methodology, data analysis, writing of the original draft. LC: data analysis, investigation, methodology, supervision, validation. IG: methodology, investigation. CR: methodology, investigation. SZ: methodology, investigation, data analysis. BB: methodology, investigation. FiC: visualization, writing review & editing. GGC: conceptualization, funding acquisition, validation, writing, review & editing. FrC: data analysis, investigation, methodology, writing of the original draft, review & editing, conceptualization, supervision, validation.

## Acknowledgments

The authors thank all members of the Neuropathophysiology lab for critical discussion of our data; Cristina Scielzo and Riccardo Pinos for setting up 3D bioprinted cultures. Image analysis was carried out at ALEMBIC, an advanced microscopy laboratory established by the San Raffaele Scientific Institute and University. We thank Hassan Marzban, Farshid Ghiyami and Azam Asemi Rad (U. of Manitoba) for helpful discussion.

## Funding

This study was supported by Ataxia Canada grant #56220 to Hassan Marzban and GGC

## Conflict of interest

The authors declare that the research was conducted in the absence of any commercial or financial relationships that could be construed as a potential conflict of interest.

---

**Supplemental Figure 1**

**No differences in astroglial marker expression between FRDA-like and control astrocytes**

**A**: Scrambled- and sh380-transduced astrocytes immunostained with an antibody detecting glial fibrillary acidic protein (GFAP, red), 7 days after transduction. Size bar: 50 μm

**B**: Percentage of GFAP+ astrocytes, over the total number of cells in the field (DAPI+ cells), analyzed by high-throughput microscopy. Each dot represents the average of a single culture well. Data, from 3 biological replicates, are expressed as mean +/- SEM; n=5801 cells for scrambled, and 5669 for sh380

**C**: Scrambled- and sh380-transduced astrocytes immunostained with an antibody detecting S100β (red), 7 days after transduction. Size bar: 50 μm

**D**: Percentage of S100β+ astrocytes, over the total number of cells in the field (DAPI+ cells), analyzed by high-throughput microscopy. Each dot represents the average of a single culture well. Data from 3 biological replicates are expressed as mean +/- SEM; n=4316 cells for scrambled, and 3352 for sh380.

## Notes

### Competing Interest Statement

The authors have declared no competing interest.

