## Supplementary figures and images for "Altered calcium responses and antioxidant properties in Friedreich’s ataxia-like cerebellar astrocytes"

### Supplemental Figures Marullo 2024

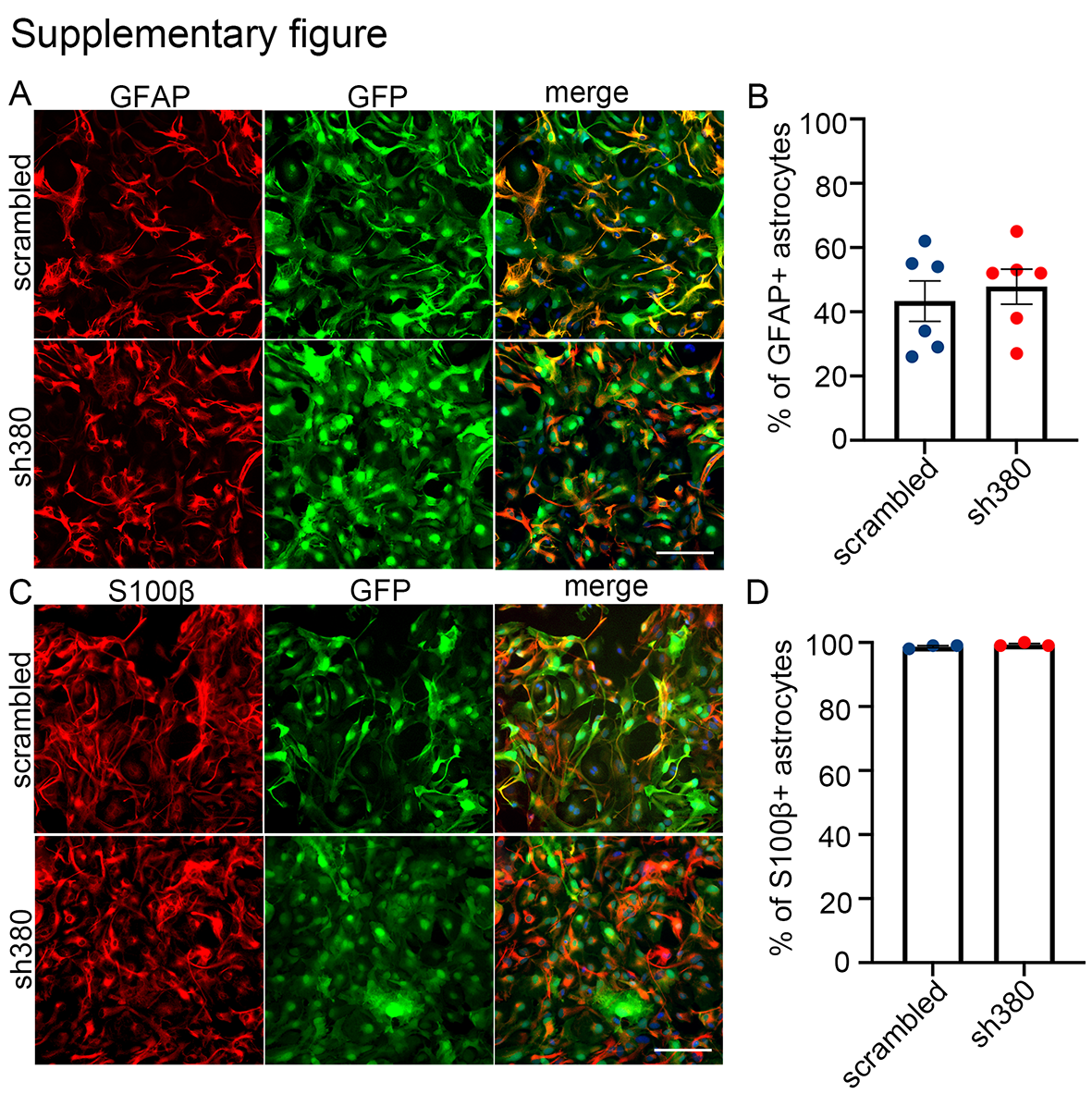
